# Disruption of hippocampal mitochondrial function underlies opioid-induced postoperative cognitive dysfunction in aged rats

**DOI:** 10.64898/2026.01.30.702943

**Authors:** Stephanie M. Muscat, Nicholas P. Deems, Bryan D. Alvarez, Michael J. Butler, Brigitte M. Gonzalez Olmo, Dominic W. Kolonay, Hannah F. Sanders, Emmanuel A. Scaria, Sabrina E. Mackey-Alfonso, Jade A. Blackwell, James W. DeMarsh, Menaz N. Bettes, Tam T. Quach, Giuseppe P. Cortese, Erica R. Glasper, Benedetta Leuner, Kedryn K. Baskin, Ruth M. Barrientos

## Abstract

Postoperative cognitive dysfunction (POCD) is a common and persistent complication in aging individuals following surgery, particularly when opioids are used for perioperative pain management. Although opioids are widely administered in the perioperative setting, the mechanisms by which they contribute to long-term cognitive impairment remain poorly understood. Here, we investigated how synaptic, neuroaxonal, and mitochondrial abnormalities contribute to long-lasting memory deficits induced by surgery and morphine, and evaluated therapeutic strategies targeting neuroinflammation and mitochondrial dysfunction. Using an aged rat model of surgery with perioperative morphine administration, we found that persistent hippocampal-dependent memory impairments were not attributable to systemic illness or gross dendritic degeneration. Instead, morphine-treated animals exhibited selective reductions in dendritic spine subtypes associated with synaptic stability, impaired late-phase long-term potentiation, and blunted experience-dependent upregulation of the AMPA receptor subunit GluA1. These synaptic alterations were accompanied by elevated circulating neurofilament light chain (Nf-L), indicating sustained neuroaxonal perturbation. Morphine treatment also produced persistent hippocampal mitochondrial dysfunction, characterized by impaired oxidative phosphorylation, reduced respiratory reserve capacity, and increased oxidative DNA damage, including mitochondrial DNA oxidation. These effects were restricted to the hippocampus and not observed in peripheral tissue. Pharmacological inhibition of TLR4 signaling at the time of surgery, which rescued the memory deficit, attenuated oxidative stress and partially restored mitochondrial function, implicating early neuroinflammatory signaling in the development of long-term mitochondrial impairment. Finally, targeted mitochondrial rejuvenation with SS-31 four weeks post-surgery robustly rescued hippocampal-dependent memory and normalized mitochondrial respiratory function despite persistently elevated DNA oxidation and Nf-L. Together these findings identify sustained hippocampal mitochondrial dysfunction as a key mechanistic substrate underlying long-term cognitive deficits following surgery and morphine exposure in aged rats, and highlight mitochondrial bioenergetics as a promising therapeutic target for POCD.

## Introduction

Gradual decline in cognition is a common feature of normal aging. However, exposure to immune challenges, such as surgery, infection, injury, or unhealthy diets, can accelerate these declines and precipitate more pronounced and persistent impairments (Barrientos, Hein, Frank, Watkins, & Maier, 2012; Barrientos et al., 2006; Godbout et al., 2005; Moller et al., 1998; Muscat et al., 2024; Muscat et al., 2021; Sparkman & Johnson, 2008; Spencer et al., 2017; Wangler et al., 2022). One clinical manifestation of this vulnerability is postoperative cognitive dysfunction (POCD), a condition characterized by persistent impairments in cognitive function, including confusion, executive dysfunction, and episodic memory loss, most commonly observed in older adults following surgery (Evered et al., 2018; Mahanna-Gabrielli et al., 2019; Moller et al., 1998; Terrando et al., 2011). Unlike postoperative delirium, which typically resolves within 7 days, POCD is defined by cognitive deficits lasting 30 days or longer after surgery (Evered et al., 2018) and is associated with an increased risk of developing Alzheimer’s disease and related dementias (Bickel, Gradinger, Kochs, & Forstl, 2008; Mahanna-Gabrielli et al., 2019).

Preclinical studies have identified neuroinflammation as a critical factor in the development of post-surgical cognitive impairments (Barrientos et al., 2012; Hovens, Schoemaker, et al., 2014; Hovens, van Leeuwen, et al., 2014; Kawano et al., 2016; Rosczyk, Sparkman, & Johnson, 2008; Wang et al., 2020). However, many of these models demonstrate only transient cognitive impairments, typically lasting 2 weeks or less, and therefore do not accurately capture the prolonged nature of POCD observed clinically. To address this limitation, our lab developed a rodent model of POCD in which multiple control conditions, including young adult animals, sham surgery, and saline-treated surgical controls, were systematically examined. These experiments demonstrated that persistent memory impairments lasting at least eight weeks emerged exclusively in aged rats that underwent surgery and received postoperative morphine (Muscat et al., 2021). Neither aging alone, surgery alone, nor morphine administration in the absence of surgery was sufficient to produce long-lasting cognitive deficits. Rather, all three factors (advanced age, surgical insult, and opioid exposure) were required to elicit enduring memory impairment. This model therefore more faithfully mirrors the chronic trajectory of POCD (Muscat et al., 2021).

Subsequent work from our laboratory revealed that these combined factors exacerbated postoperative cognitive deficits through enhanced neuroinflammatory signaling, specifically via activation of the innate immune receptor toll-like receptor 4 (TLR4) (Muscat et al., 2023). This model has strong clinical relevance, as nearly 90% of surgical patients are prescribed morphine or other opioids for postoperative pain management (Aubrun, Mazoit, & Riou, 2012; Li et al., 2018; Lindenhovius et al., 2009; Thiels et al., 2017), older adults undergo more surgical procedures than any other age group (Fowler, Abbott, Prowle, & Pearse, 2019), and they are at heightened risk for POCD (Bickel et al., 2008; Leighton et al., 2022; Wacker, Nunes, Cabrita, & Forlenza, 2006). Interestingly, although neuroinflammation plays a key role in initiating postoperative cognitive decline, memory impairments persist well beyond the resolution of the neuroinflammatory response in this model (Muscat et al., 2023; Muscat et al., 2021), suggesting that additional downstream mechanisms sustain long-lasting cognitive dysfunction.

The current study aimed to identify these downstream mechanisms contributing to persistent POCD following surgery and morphine exposure in aged rats. We hypothesized that this condition would be associated with impaired synaptic plasticity, structural abnormalities in dendritic spines, and altered expression of glutamate receptors, processes that are critical for normal long-term memory function (Bosch & Hayashi, 2012; Collingridge & Abraham, 2022). We further investigated how early neuroinflammatory signaling might drive these neuronal alterations. Because dendritic spine remodeling and synaptic transmission are highly energy-dependent processes (Camandola & Mattson, 2017), and neuroinflammation is known to elevate reactive oxygen species (ROS) that impair mitochondrial function and energy metabolism (Chen & Zweier, 2014; Lin et al., 2022), we hypothesized that surgery and morphine would lead to sustained oxidative stress and mitochondrial dysfunction in the hippocampus. Finally, to determine whether mitochondrial dysfunction represents a viable therapeutic target once POCD is established, we tested whether pharmacological mitochondrial rejuvenation with the mitochondrial-targeted peptide SS-31 could reverse cognitive and cellular deficits when administered four weeks after surgery.

Taken together, these experiments were designed to define a mechanistic pathway linking early postoperative neuroinflammation to long-lasting mitochondrial, synaptic, and cognitive dysfunction, and to evaluate whether restoring mitochondrial bioenergetics is sufficient to rescue memory in aged animals with POCD.

## Materials and Methods

### Experimental design

This study was comprised of six separate experiments that, for ease of reading, will be briefly summarized here. Specific methodological details (such as dose, route of administration, etc.) will be described in the appropriate subsections below. Based on our prior studies described earlier (Muscat et al., 2023; Muscat et al., 2021), the present study focused exclusively on aged rats that underwent surgery, allowing us to interrogate mechanisms underlying persistent POCD without redundancy from experimental conditions previously shown to be insufficient. Thus, young adult rats and sham-operated controls were not included in these studies.

In **Experiment 1**, we aimed to determine if the combination of aging, surgery, and morphine was associated with neuronal structural pathology at the 30-day post-surgical timepoint. Rats received the surgical procedure (laparotomy) on day 1 followed by 6 days of morphine treatment. On day 30, they underwent a contextual learning experience (they were exposed to a novel context) and were euthanized 2 hours later, when memory consolidation processes are known to be peaked (Barrientos et al., 2004). Brains were extracted, processed, sliced, and Golgi-Cox stained for visualization of neuronal morphology. Dendritic complexity and dendritic spines were quantified and analyzed.

In light of Experiment 1 findings, which showed that mature dendritic spines were diminished with surgery and morphine, **Experiment 2** determined the functional consequence of this reduction and assessed synaptic plasticity. Aged rats received laparotomy and morphine just as in Experiment 1, and on day 30, they were euthanized, brains were extracted, and hippocampal slices were prepared to measure long-term potentiation (LTP).

After finding that synaptic plasticity was severely inhibited at 30 days post-surgery and morphine, **Experiment 3** assessed protein expression of neurofilament light chain (Nf-L) in the circulation, and various glutamate receptor subunits in hippocampal synaptoneurosomes. Aged rats received laparotomy and morphine just as in Experiment 1, and on day 30, they were euthanized 2 hours after a learning experience, and serum and brains were collected. Nf-L was measured via ELISA. Synaptoneurosomes were isolated from the hippocampus and protein expression of various glutamate receptor subunits were measured via western blot analysis.

**Experiment 4** aimed to determine the extent to which surgery and morphine would induce DNA oxidation and mitochondrial dysfunction. Aged rats received laparotomy and morphine just as in Experiment 1, and on day 30, they were euthanized 2 hours following learning exposure and hippocampus from each brain hemisphere were dissected. One hemisphere was frozen and later processed for DNA oxidation analysis via ELISA. From the other hemisphere, mitochondria were freshly isolated to assess oxygen consumption rate in two separate Seahorse assays.

In **Experiment 5**, we aimed to determine the extent to which early neuroinflammation, caused by combined surgery and morphine, directly contributes to the mitochondrial dysfunction observed at the advanced stages (Day 30) of POCD. Thus, we pharmacologically blocked TLR4 signaling in the brain at the time of surgery with LPS-RS Ultrapure, the specific TLR4 receptor antagonist. Immediately following this injection, aged rats received laparotomy and morphine treatment as in Experiment 1. On day 30, they were euthanized 2 hours after learning and hippocampi were processed to measure DNA oxidation and mitochondrial respiration, as in Experiment 4.

Given mitochondrial dysfunction consistently emerged as a prominent feature of the hippocampus 30 days after surgery across prior experiments, **Experiment 6**, examined whether treatment with SS-31, a drug known to rejuvenate mitochondrial structure and function, could reverse surgery and morphine-induced deficits in mitochondrial respiration and memory. Aged rats received laparotomy and morphine just as in Experiment 1, and at 30 days post-surgery, they were administered SS-31 every other day (for a total of 3 doses) prior to receiving a learning experience and long-term memory was assessed four days later. Rats were then euthanized and hippocampi were processed to measure DNA oxidation, Nf-L, and mitochondrial respiration, as in Experiment 4.

### Animals

Aged (22-24 months old), male F344xBN F1 rats obtained from the National Institute on Aging Rodent Colony managed by Charles River were used. Unfortunately, female rats of this strain and age, exclusively available through the NIA colony, were not available at the time these studies were conducted. Our previous work using this age and strain of rats has indicated that unchallenged rats exhibit no significant differences in memory performance compared to young adult (3 months old) controls. Young adult rats were not used in this study based on our previous work that established they are not impaired in this model (Muscat et al., 2021).

Condition-matched rats were housed 2 to a cage (52 L x 20 W x 21 H, cm). The animal colony was maintained at 22±2°C on a 12h light/dark cycle (lights on at 700 h). Rats were allowed *ad libitum* access to food and water. Animals were given at least 1 week to acclimate to colony conditions prior to experimentation. All experiments were conducted in accordance with protocols approved by the Ohio State University Animal Care and Use Committee. Every effort was made to minimize the number of animals used and their suffering.

### Surgery

Laparotomy (exploratory abdominal surgery) was performed using aseptic procedures under isoflurane anesthesia, as described previously (Barrientos et al., 2012; Muscat et al., 2024; Muscat et al., 2023; Muscat et al., 2021). Briefly, the abdominal region was shaved and cleaned with alternating 70% ethanol and surgical scrub. Approximately 0.5 cm below the lower right rib, a 3 cm incision was made, revealing the peritoneal cavity. While wearing sterile gloves, the musculature was outstretched to further open the incision. Then, approximately 10 cm of the small intestines were exteriorized and vigorously manipulated for 30 s, after which they were returned into the peritoneal cavity. Sterile chromic gut sutures (3-0) were used to suture the peritoneal lining and abdominal muscle in two layers; the skin was closed with surgical staples.

The wound was then dressed with triple antibiotic ointment to prevent infection. Each procedure lasted ∼20 min. Based on previous work demonstrating sham-operated rats are unimpaired at any and all time points, sham surgery controls were not included in this study (Barrientos et al., 2012; Muscat et al., 2021). Because the present study assessed the impact of an analgesic (morphine), control animals did not receive any analgesic treatment to avoid potential confounds. All subjects were assessed twice daily for the first week after surgery, and then twice weekly thereafter, to monitor surgical recovery.

### Drugs and administration procedures

Morphine was gifted by the NIDA drug repository and administered i.p. at a dose of 2 mg/kg/ml twice daily (∼900 h and ∼1700 h) for seven days, based on prior studies (Muscat et al., 2023; Muscat et al., 2021). The human morphine dose equivalency to the rat dose used is 45 mg/day (based on calculations suggested in (Reagan-Shaw, Nihal, & Ahmad, 2008) and was chosen based on the recommended dose for opioid-naïve patients (MD Anderson Cancer Center, 2018). Morphine is reported as a free base concentration and was diluted in sterile saline (0.9%). An equivolume of sterile saline was administered to control animals.

For Experiment 5, we used lipopolysaccharide from the photosynthetic bacterium *Rhodobacter sphaeroides* (LPS-RS Ultrapure, InvivoGen) as a competitive antagonist of TLR4 via its binding to the MD-2 coreceptor binding site (Coats, Pham, Bainbridge, Reife, & Darveau, 2005; Visintin, Halmen, Latz, Monks, & Golenbock, 2005). LPS-RS Ultrapure (hereafter referred to simply as LPS-RS) is a selective TLR4 inhibitor with no activity at TLR2 (Hirschfeld, Ma, Weis, Vogel, & Weis, 2000; Vallance et al., 2019). It was injected intracisterna magna (icm) at a dose of 50 μg/5 μL, based on (Muscat et al., 2023). Icm administration was utilized to directly target the CNS in a minimally invasive manner, to reduce additional inflammation that may be caused by stereotaxic surgery. Under isoflurane anesthesia, the dorsal aspect of the skull was shaved and then cleaned with 70% ethanol. A 27-guage needle attached via PE50 tubing to a 25 μL Hamilton syringe was inserted into the cisterna magna. Entry into the cistern was verified by drawing up ∼2 μL of clear cerebral spinal fluid. The cerebral spinal fluid was then gently pushed back in and 5 μL total volume of LPS-RS was subsequently administered over 30 s. The entire procedure was completed in ∼3 min. Vehicle control animals received an equal volume of sterile water.

For Experiment 6, the compound elamipretide, also known as SS-31, was injected 30 days after surgery at a dose of 5mg/kg, i.p. every other day for three total administrations. The last dose corresponded to the day before the learning session. Dose and length of dosing was based on several published studies (Avery et al., 2024; Czigler et al., 2019; Miyamoto et al., 2020; Nickel et al., 2022; Reddy, Manczak, & Kandimalla, 2017; Zhang et al., 2017) and our own pilot data.

### Golgi-Cox Staining, slicing & imaging

Following rapid decapitation, brains were removed and rinsed briefly in distilled water to remove excess blood, then immersed in Gogi-Cox impregnation solution according to manufacture instructions (F&D Neurotech). Briefly, brains were placed in specimen jars containing 10 mL of freshly prepared working solution and stored in the dark at room temperature for 2 weeks, with solution changes performed according to manufacturer’s recommendations. After impregnation, brains were transferred to cryoprotectant solution provided by the manufacturer and stored in the dark at 4°C for 3 days. Tissue was then rapidly frozen and sectioned coronally on a cryostat at −20° C at a thickness of 150 um. Sections were mounted on Superfrost Plus glass slides coated with poly-L-lysine, and allowed to dry overnight at room temperature in the dark. Slides were subsequently developed and stained according to manufacturer’s protocol. Following staining, sections were then dehydrated in a series of graded ethanols (50%, 75%, 95%, and 100%), cleared in CitriSolv, and cover-slipped using a permanent mounting medium.

Images were obtained by optical microscopy (Nikon 90i-Eclipse Confocal-Brightfield). Dendrites from hippocampal neurons in the dorsal CA1 region were imaged at 40x. Reconstruct was used to trace and generate 3D reconstructions of dendrites and spines. Sholl analysis was performed using the NIH Image J Neuroanatomy plugin, and area of dendritic arbor, branch tips, stems, bifurcations, and length were used to gauge complexity of imaged neurons. Dendritic spines were measured from 10um segments of second-order branches using a 100x oil objective.

### Dendritic Spine Analysis

NIS Elements was used to automate dendritic spine classification using the machine learning plugin (NIS AI). Because NIS Elements analysis cannot determine the orientation of images or objects, it relies on its internal length metric (the longest axis of an object) to measure object length and the perpendicular minor axis to estimate width. After optimizing and comparing automated measurements with manual measurements of spine length and width, we found that length measurements were more accurate, largely due to the elongated and irregular morphology of dendritic spines.

To enable accurate spine classification, NIS AI was trained to generate two distinct 2D object overlays using eight dendrites from the same dataset that was subsequently analyzed (Supplementary Figure S1): one overlay capturing spine heads (head.oai) and a second overlay encompassing the entire spine (spine.oai). This approach allowed both spine head dimensions and total spine dimensions to be derived from the NIS length metric, which measures the longest dimension of an object regardless of its orientation. As a result, both the length of the entire spine and the effective width were objectively measured using the length-based formula for every spine. Dendritic spines were classified into morphological subtype using a quantitative adaptation of the unbiased classification scheme described previously (Risher, Ustunkaya, Singh Alvarado, & Eroglu, 2014). Classification was based on direct morphometric measurements of spine length and head width obtained from NIS Elements, from which length-to-width ratio (LWR) was calculated for each spine. Rather than relying on subjective visual assessment, spines were assigned to mutually exclusive morphological categories using predefined numerical thresholds consistent with those previously proposed (Risher et al., 2014). Specifically, filopodia were defined as protrusions with length > 2.0 μm, mushroom spines were identified by a head width > 0.6 μm; long thin spines had a length > 1.0 μm without meeting mushroom criteria, thin spines were defined by an LWR > 1.0 and stubby spines were classified by an LWR < 1.0. Classification followed a hierarchical decision structure to ensure that each spine was assigned to a single morphological category. In addition, the physical length of the imaged regions in each image was recorded, enabling normalization of spine measurements on a per-micron basis (Supplementary Figure S2).

Because images were not originally acquired with this analysis pipeline in mind, each image required manual removal of 2D overlays generated on non-target dendrites. To ensure accurate pairing of spine heads with their corresponding spine bodies, only spine bodies that overlapped with a detached spine head were included in the analysis. Although all images experienced a small degree of attrition, the application of a consistent and objective machine-learning-based classification mitigates this limitation. Lastly, both training datasets (head.oai and spine.oai) were trained through 3,000 iterations. The final training loss was 0.00474 for the heads.oai model and 0.00363 for spine.oai model, both well below the maximum recommended training loss threshold of 0.015, in accordance with Nikon’s NIS AI technical support.

### Long-term potentiation

Late-phase long-term potentiation (L-LTP) in a hippocampal slice preparation and extracellular recording procedure was utilized to assess synaptic plasticity as we have done previously (Chapman, Barrientos, Ahrendsen, Maier, & Patterson, 2010; Gonzalez Olmo et al., 2023; Tanaka, Cortese, Barrientos, Maier, & Patterson, 2018). Four weeks post-surgery, rats were rapidly decapitated without anesthesia prior to tissue collection. Brains were carefully extracted from the skull and placed in an ice-cold slicing solution (in mM: 250 Sucrose, 24 NaHCO3, 25 Glucose, 2.5 KCl, 1.25 NaH2PO4, 1.5 MgSO4, 2 CaCl2; pH adjusted to 7.3-7.4) for approximately 20 minutes prior to slicing. Transverse hippocampal slices were collected using a vibratome. The brain was transferred to a cold Petri-dish with the same ice-cold slicing solution. To obtain hippocampal sections, cerebellum and a small part of the prefrontal cortex was cut off. The remaining block was cut along the midline into two equal hemispheres using a scalpel blade. Both hemispheres were mounted to the specimen holder with superglue and a supporting piece of agar behind the brain, away from the side of the vibratome, to provide structural support during slicing. The vibratome was set up by filling the buffer tray with ice-cold slicing solution, ice bath tray with ice to keep the buffer tray cold during slicing, and adjusting the desired thickness to 400 μm and cutting speed to 0.06 mm/s. Using a transfer pipette, slices were transferred to the slice incubation chamber that was placed in a water bath to be maintained at 28°C and perfused with oxygenated aCSF (in mM: 124.0 NaCl, 4.4 KCl, 26.0 NaHCO3, 1.0 NaH2PO4, 2.5 CaCl2, 1.3 MgSO4, 10 Glucose). After slicing, the chamber was removed from the water bath and slices were permitted to recover from the mechanical shock of slicing for at least 90 min at room temperature before recording. Field excitatory postsynaptic potentials (fEPSPs) were recorded from Schaffer collateral–CA1 synapses by placing both stimulating and recording electrodes in the stratum radiatum. All stimuli were delivered at intensities that evoked fEPSP peak amplitude approximately 35%-50% of the maximum response recorded during I/O measurements in each slice. Percentage of facilitation of PPF was calculated from the ratio of the second fEPSP peak amplitude to the first fEPSP peak amplitude, shown at intervals ranging from 50 to 250 ms. Test stimuli were delivered once every minute, and test responses were recorded for 30 min before beginning the experiment to assure stability of the response. The same stimulus intensity was used for LTP induction and to evoke test responses. Theta-burst stimulation (TBS) protocol (12 bursts of 4 pulses at 100 Hz, delivered 200 ms apart) was used to induce late-phase LTP (L-LTP), a correlate of long-term memory (Abraham, 2003; Bramham & Messaoudi, 2005; Frey & Morris, 1997), and recording lasted 60 minutes. fEPSP mean peak amplitude was used as a measure of synaptic activity. The theta-burst protocol, recognized as a more robust LTP induction paradigm, is designed to mimic the burst firing of CA1 pyramidal cells at theta frequency recorded in vivo from awake animals during spatial exploration (Abraham & Huggett, 1997; Larson, Wong, & Lynch, 1986; O’keefe, 2007). This protocol is considered to be more naturalistic and has proven to be a sensitive indicator of alterations in mnemonic processes associated with aging (Chapman et al., 2010; Gonzalez Olmo et al., 2023; Tanaka et al., 2018)

### Neurofilament Light-Chain Assay

Neurofilament light-chain (Nf-L), a cytoskeletal protein enriched in large-caliber axons and is widely used as a sensitive marker of axonal injury, integrity, and neuroaxonal turnover (Gafson et al., 2020; Pekny et al., 2021), was measured in serum using MSD’s R-PLEX Human Neurofilament L Assay (Catalog No K1517XR-2) according to the manufacturer’s instructions. Samples were diluted two-fold with diluent 11. The calibration curve was constructed using eight calibrators with concentrations ranging from 0 to 50,000 ng/L. Nf-L concentrations were calculated from calibrator signals using a 1/Y^2^-weighted four-parameter logistic (4PL) curve fit.

### Synaptoneurosome preparation

All tissue was collected 4 weeks post-surgery. On the day of tissue collection, rats were repeatedly exposed to a novel context to engage the hippocampus, and were euthanized 2 hours later, when memory consolidation processes are known to be active (Barrientos, O’Reilly, & Rudy, 2002; Barrientos et al., 2004). Four weeks post-surgery was selected because of the following: (1) all of the animals have completely recovered from the surgery; (2) the aging, but not the young rats exhibit significant impairments in long-term memory (Muscat et al., 2021); and (3) levels of proinflammatory cytokines in the hippocampus are no longer significantly elevated in the aging rats (Muscat et al., 2023; Muscat et al., 2021). Rats underwent rapid decapitation, and hippocampi were extracted. Synaptoneurosomes were isolated using well-established methods (Cortese, Barrientos, Maier, & Patterson, 2011). Tissue was placed in 2 mL of SynPer buffer (Thermo) with protease and phosphatase inhibitors and homogenized using a glass tissue grinder and pestle. Nuclear material and unbroken cells were removed by centrifugation at 1200 x g for 10 min at 4° C. The remaining supernatant was centrifuged at 15,000 x g for 20 min at 4° C, yielding an S2 cytosolic fraction and a P2 pellet containing the synaptoneurosomal fraction comprised of both presynaptic and postsynaptic material. The P2 synaptoneurosomal pellet was resuspended in 500 ul of Syn-Per reagent. The P2 fraction obtained using this protocol is enriched for perisynaptic components including presynaptic and postsynaptic proteins, terminal mitochondria and cytoplasm and synaptic vesicles (Booth & Clark, 1978; Whittaker, 1993). Synaptic enrichment of the P2 fraction was confirmed by the presence of synaptophysin and postsynaptic density 95 (PSD95) in the P2 fraction and their absence from the S2 fraction by western blot (data not shown). Samples were stored in 5% (v/v) DMSO at −80° C until western blots were completed.

### Western blots

The P2 fraction of synaptoneurosomes were used to measure protein expression of glutamate receptor subunits. Bradford protein assays were performed on all samples to determine total protein concentrations. Samples were divided into 10 μL aliquots and frozen at −80°C until western blots were performed. An equal amount of total protein (50 μg) from each sample was loaded into each lane. The NuPAGE Bis-Tris (10 well, 4-12%, 1.5 mm) gel electrophoresis system was used under reducing conditions (Life Technologies). The iBlot dry-blotting system (Life Technologies) was used to electrophoretically transfer gels to nitrocellulose membranes. Odyssey TBS blocking solution (Li-Cor) was used to prevent nonspecific protein binding (1 hr at RT). Primary antibodies, in Odyssey blocking solution containing 0.2% Tween20 (overnight at 4°C), were: SYP (1:1,000; sc-17750; Santa Cruz), PSD95 (1:1,000; 2507S; Cell Signaling), GluA1 (1:1000; 13185S; Cell Signaling) GluA2/3 (1:1000, 07-598; Millipore Sigma), GluN1 (1:500; MAB1586; Millipore Sigma), GluN2A (1:500; 07-632; Millipore Sigma) and GluN2B (1:1000; ab254356; Abcam),

### Seahorse Mitochondrial Respiration Assays

Oxygen consumption rates (OCR) were assessed in mitochondria isolated from fresh hippocampal tissue with two complementary assays using the Seahorse XFe96 Bio-analyzer as we have done previously (Butler et al., 2026). The electron coupling (EC) assay measures coupling between the electron transport chain (ETC) and oxidative phosphorylation, and the electron flow (EF) assay determines the sequential flow of electrons through the different ETC complexes independent of the proton gradient/membrane potential (Rogers et al., 2011). Following rapid decapitation, one hemisphere of the hippocampus was dissected and placed in cold mitochondrial isolation buffer and homogenized in a glass tube using an electric rotator. Homogenized tissue was centrifuged at 800 x g for 10 min at 4° C. The lipid layer was aspirated, and the supernatant was passed through cheese cloth into a clean tube. This was then centrifuged at 8000 x g for 10 min at 4° C. Supernatants were aspirated and the remaining mitochondrial pellet was resuspended in MSHE buffer with BSA. To obtain semi-pure mitochondrial samples, two additional rounds of centrifugation at 8000 x g for 10 min at 4° C were completed and mitochondria were resuspended with MSHE buffer without BSA.

A Bradford assay was conducted on the resuspended mitochondria to estimate the number of mitochondria based on protein concentration. For the EC assay, 2 μg of hippocampal mitochondria in 25 μL EC assay buffer were plated in a 96-well Seahorse microplate. The plate was centrifuged at 2,000 x g for 20 minutes at 4° C, overlaid with 155 μL of the EC assay buffer containing excess succinate (10mM) and rotenone (2 μM), and incubated at 37° C in a CO_2_-free environment for 8-10 minutes prior to the assay. Three OCR measurements were taken at each phase of the assay including at baseline and following the sequential addition of ADP (400 μM well concentration), oligomycin (3 μM well concentration), carbonyl cyanide-4 (trifluromethoxy) phenylhydrazone (FCCP; 8 μM well concentration), and a combination of rotenone and antimycin A (4 μM well concentration). For the EF assay, 2μg isolated hippocampal mitochondria were plated in 25 μL EF assay buffer containing excess substrates pyruvate (10 mM), malate (2 mM), and FCCP (4 μM). After centrifugation and addition of 155 μL of the EF assay buffer, the assay was run with sequential addition of rotenone (2 μM well concentration), succinate (15 mM well concentration), antimycin A (4 μM well concentration), and TMPD (N, N, N′, N′-tetramethyl-p-phenylenediamine; 150 μM well concentration). To determine whether mitochondrial dysfunction was limited to brain tissue or if it was also disrupted in peripheral tissues, we measured OCR in liver. Because liver tissues were frozen, inducing mitochondrial uncoupling, a modified EF assay was performed. 7μg isolated liver mitochondria were plated in 25 μL EF assay buffer containing excess substrates pyruvate (10 mM), malate (2 mM), and FCCP (4 μM). After centrifugation and addition of 155 μL of the EF assay buffer, the assay was run with sequential addion of succinate (10 mM well concentration), antimycin A (4 μM well concentration), and TMPD (150 μM well concentration). To analyze data, OCR area under the curve (AUC) of each phase of each assay was calculated for each biological replicate and then groups differences were analyzed with a Welch’s t-test (Exp. 4 and 6) or one-way ANOVA (Exp. 5).

### DNA Oxidation

Oxidative stress was assessed using a DNA/RNA Oxidative Damage (High Sensitivity) ELISA Kit (Catalog No. 589320, Cayman Chemical), which is a competitive assay that measures 8-hydroxyguanosine (8-OHG), 8-hydroxy-2’-deoxyguanosine (8-OHdG), and 8-hydroxyguanine. Protein sonicates from the hippocampus and prefrontal cortex were diluted 2-fold (i.e. 30ul of sample: 30ul of ELISA buffer) for the assay. The ELISA was performed according to manufacturer’s instructions and absorbance was measured at 450nm. Each sample was normalized to total amount of protein (as measure by Bradford assay) and analyzed as pg/ug total protein.

### Contextual fear conditioning

Memory function was assessed using a modified version of contextual fear conditioning known as the context pre-exposure facilitation effect paradigm. This assay was chosen because it is highly and specifically dependent on the hippocampus as it allows the rats to learn the context incidentally, independent of its association with the aversive shock (Fanselow, 1990; Rudy, Barrientos, & O’Reilly, 2002). Additionally, we have validated its use in detecting hippocampal memory deficits following an array of inflammatory insults (Barrientos et al., 2006; Muscat et al., 2023; Muscat et al., 2021; Sobesky et al., 2014), and have established that surgery in aged rats impairs hippocampal memory but does impact amygdalar-dependent memory (Barrientos et al., 2012). Rats were conditioned using the Coulbourn Instruments Habitest Modular System. Each conditioning chamber (12” W x 10” D x 12” H) had two solid metal walls and two walls made of Plexiglas. An audio speaker and a house light were mounted on the ceiling. A foot shock could be delivered through a removable grid shock floor. The rods were wired to a shock generator for eight unique shock outputs. These conditioning chambers were placed inside an isolation cubicle (23” W x 20” D x 24” H). Before each animal was conditioned or tested, the chambers were cleaned with water and 70% ethanol.

Briefly, this task is comprised of three components: a pre-exposure phase, an immediate shock phase, and a memory retrieval phase. During the pre-exposure phase, rats were transported in a black bucket from their home cage to the conditioning context, where they were allowed to freely explore. This was repeated six times (rats remained in the conditioning context for 5 min on the first exposure and for 40 sec on each of the five subsequent exposures, with an approximately 40 sec interval in their home cage between each subsequent exposure) in order to establish an association between the black bucket and activation of the conjunctive representation of the conditioning context as described previously (Rudy et al., 2002). During the first, 5 min exposure, general locomotion was assessed to account for any confounding motor disturbances or generalized fear. 3 days later, the second phase, immediate shock phase was carried out. Rats were transported from their home cage to the conditioned context in the same black bucket as before. Immediately upon being placed in the conditioned context, rats received one 2 sec, 1.5 mA foot shock. They were then removed from the conditioned context and transported back to their home cage. During the immediate shock phase, the rats’ time in the conditioned context never exceeded 10 sec. 24 hours after the immediate shock phase (4 days after pre-exposure), rats were tested for memory of the conditioned context. During this testing phase, rats were again transported from their home cage to the conditioned context in the black bucket. They were then observed and scored for freezing behavior; freezing is a rat’s dominant fear response and is characterized by a complete suppression of behavior, including immobility, shallow breathing, and autonomic changes such as increased heart rate and piloerection (Fanselow and Lester, 1988). In this study, freezing was defined as the absence of all visible movement, except for respiration. Rats were scored every 10 seconds while in the chamber for 6 minutes. 2 hours later, locomotion and generalized fear were again assessed by placing the rats in a new, novel/neutral context (a round plastic chamber with wire walls and a solid floor with corn cob bedding) for another 6 minutes and observing for freezing behavior. Scoring was carried out manually in real-time by three observers blind to treatment conditions; scores were averaged, and inter-rater reliability exceeded 97% for all studies.

### Statistical analyses

For all experiments, n=6-8 rats/group were utilized, based on statistical power established by our previous work (Barrientos et al., 2012; Muscat et al., 2021; Muscat et al., 2023). Statistical analyses were performed using Prism v.10.3 software. Two-way ANOVAs, one-way ANOVAs, or Welch’s t-tests were used where appropriate, based on experimental design. Following significant main effects or interactions, Tukey’s *post hoc* tests were conducted to assess for pairwise differences between groups. Statistical significance for all tests was set at alpha=0.05.

## Results

### Experiment 1

#### Body Weight

Body weight was monitored from the day of surgery through four weeks post-surgery in saline- and morphine-treated rats. A two-way ANOVA revealed a significant main effect of time on body weight (F(1.54, 24.61) = 70.16, p < 0.0001) as well as a significant treatment x time interaction (F(1.54, 24.61) = 3.88, p < 0.05; **Fig. 1A**). Both groups experienced a modest but significant reduction in body weight (∼ 2-5%) from days 2-14 following surgery relative to pre-surgery values. Morphine-treated rats had significantly lower body weight than saline-treated rats on post-surgery days 5-7 only (p < 0.05). Body weights in both groups returned to pre-surgery levels by week 3.

**Figure 1.**
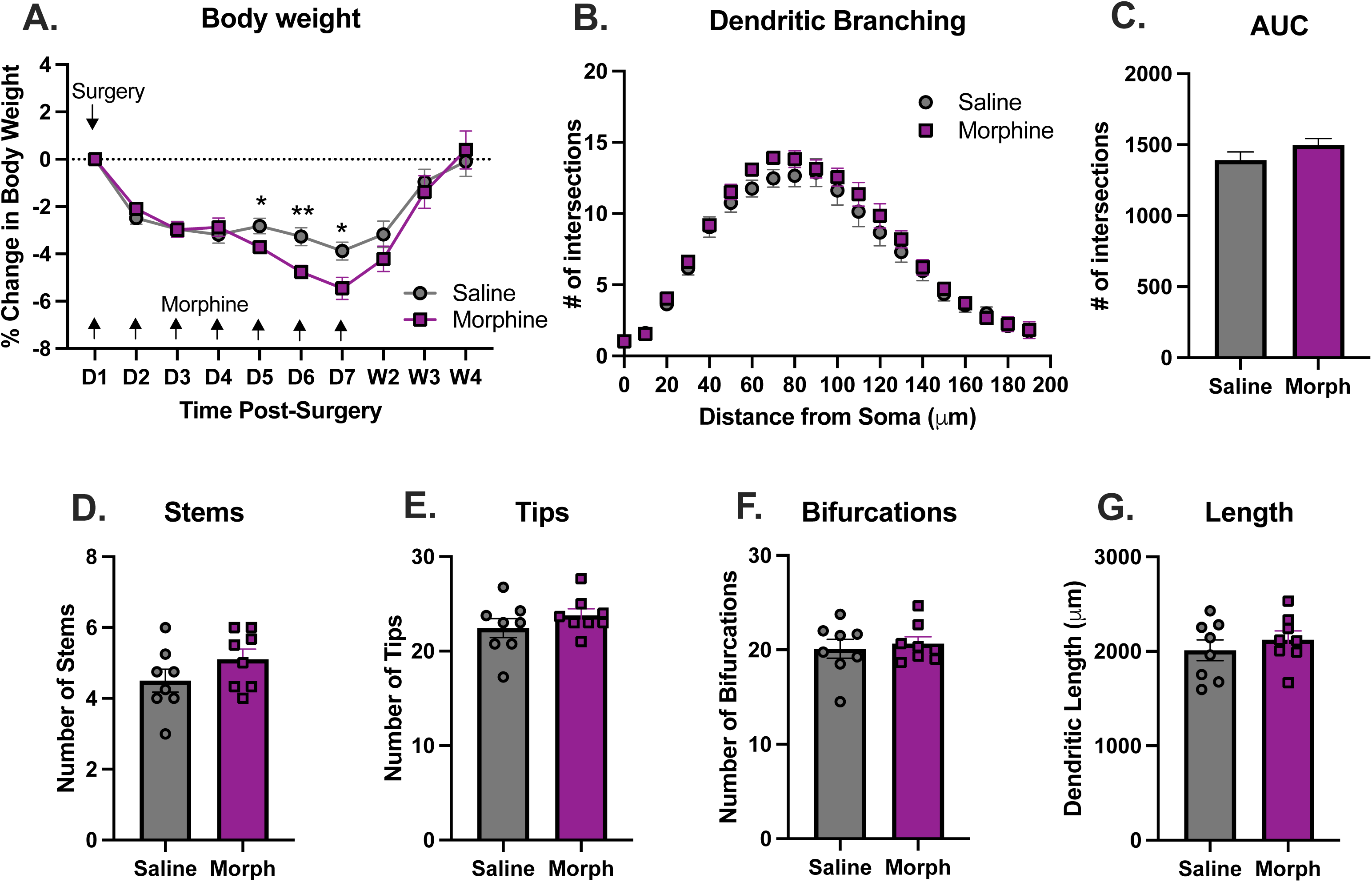
Body weight recovery and dendritic arborization. (A) Percent change in body weight relative to pre-surgery baseline across the post-surgical period for saline- and morphine-treated rats. Arrows indicate days of morphine administration. *p < 0.05, **p < 0.01. (B) Sholl analysis of CA1 pyramidal neuron dendritic branching showing the number of dendritic intersections plotted as a function of distance from the soma for saline- and morphine-treated groups. (C) Area under the curve (AUC) values calculated from Sholl profiles for each treatment group. (D-G) Quantification of dendritic architecture parameters, including number of primary stems (D), number of dendritic tips (E), number of bifurcations (F), and total dendritic length (G) in saline-and morphine-treated animals. Data are presented as mean ± SEM with individual data points shown.

#### Dendritic branching

Dendritic complexity was quantified using Sholl analysis. Morphine treatment did not significantly alter overall dendritic arborization relative to saline controls. To analyze number of intersections across distance, we calculated the area under the Sholl curve (AUC) for each rat. Saline treated rats’ AUC (1392 ± 57.76) was not significantly different than that of morphine treated rats (1498 ± 45.54), t(278) = 1.44, p > 0.05 (**Fig. 1B-C**). To summarize global differences in branching number of stems, tips, and bifurcations, and dendritic length were all calculated and determined to not differ significantly compared to saline-treated controls (p > 0.05; **Fig. 1D-G**).

#### Dendritic spine density

Spine density quantification was carried out for five different spine types (filopodia, long/thin, thin, stubby, and mushroom) in the two dendritic branches (apical and basal) of pyramidal neurons in the CA1 region of the hippocampus. In the apical branch, there were significantly fewer mushroom spines in the morphine-treated group compared to saline controls (t(16) = 4.233, p < 0.001), but density of all other spines were not significantly different (filopodia (t(16) = 0.612, p > 0.05), long/thin spines (t(16) = 0.724, p > 0.05), thin spines (t(15) =1.879, p > 0.05, stubby spines (t(16) = 0.0459, p > 0.05); **Fig. 2A-E**). In the basal branch, both mushroom (t(15) = 3.180, p < 0.01) and thin spines (t(15) = 2.721, p < 0.05) were significantly reduced in the morphine-treated group, but density of all other spine types were not different between the groups (filopodia (t(15) = 0.094, p > 0.05), long/thin (t(15) = 0.0647, p > 0.05, stubby (t(15)= 0.171, p > 0.05); **Fig. 2F-J**). Collectively, these data indicate a selective reduction in mushroom spines, a mature and stable spine subtype linked to enduring synaptic connections, in the later stages of POCD induced by surgery and morphine.

**Figure 2.**
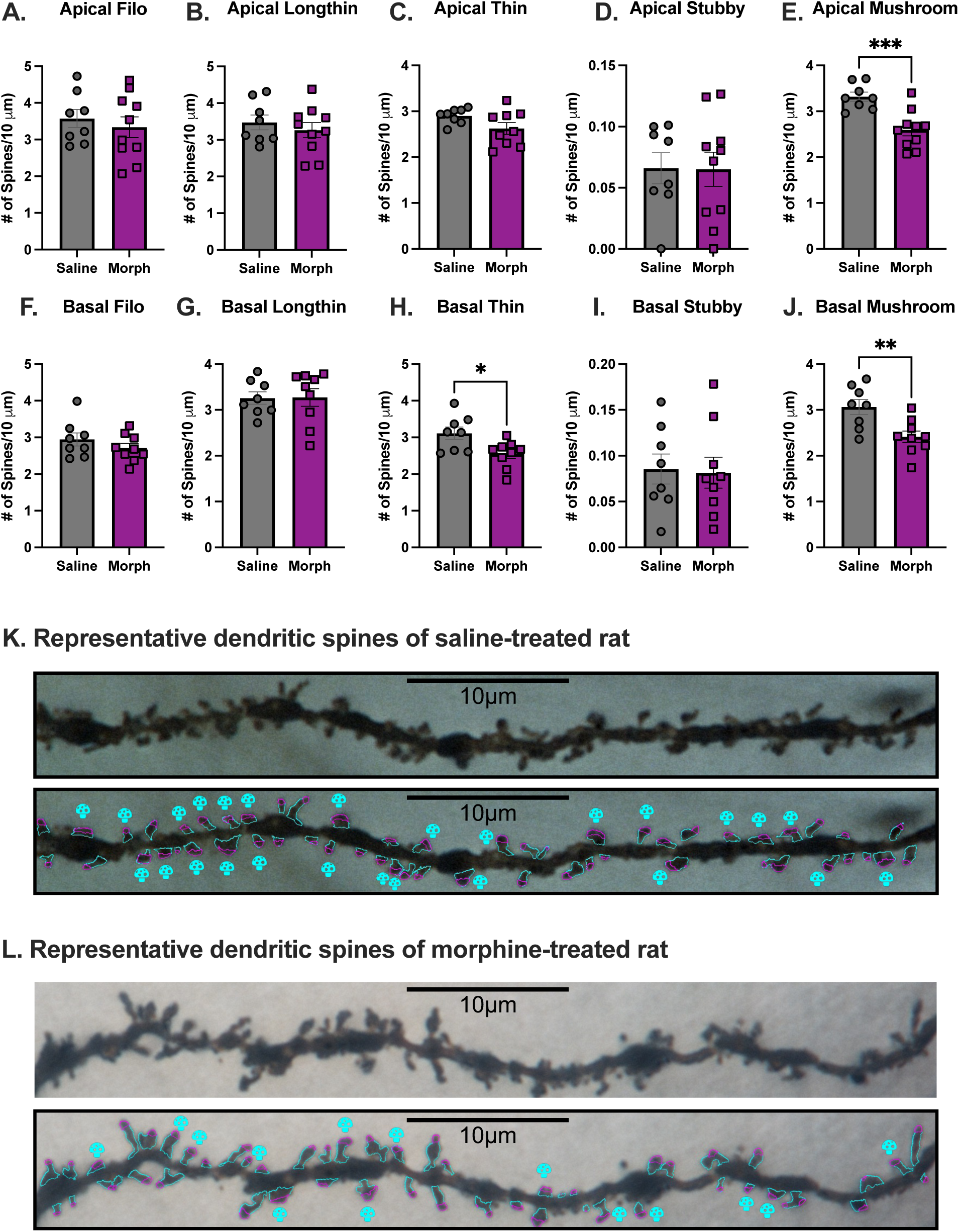
Dendritic spine density and morphological classification in CA1 pyramidal neurons. (A-E) Quantification of *apical* dendritic spine density by morphological subtype, including filopodia (A), long thin (B), thin (C), stubby (D), and mushroom (E) spines, expressed as number of spines per 10 µm of dendrite for saline- and morphine-treated rats.(F-J) Quantification of *basal* dendritic spine density by morphological subtype, including filopodia (F), long thin (G), thin (H), stubby (I), and mushroom (J) spines, expressed as number of spines per 10 µm of dendrite for each treatment group. Data are presented as mean ± SEM with individual data points shown. *p < 0.05, **p < 0.01, ***p < 0.001. (K-L) Representative image of dendritic spines from (K) a saline-treated or (L) a morphine-treated rat, shown with automated spine identification overlays. Mushroom spines are marked with a blue- colored mushroom. Scale bars = 10 µm.

### Experiment 2

#### Long-term Potentiation

Based on previously published data showing that short-term memory was intact and only long-term memory was impaired (Muscat et al., 2021), here we investigated L-LTP. At baseline, there was no significant difference between conditions (F_(1,27)_ = 0.019, p > 0.05) and these values did not differ across the 30 minutes of baseline measurements (F_(2.89, 70,22)_ = 0.541, p > 0.05). In contrast, following theta burst stimulation (TBS), saline-treated animals exhibited robust firing whereas morphine treatment severely diminished firing (F_(1,27)_ = 18.70, p < 0.001), and these recordings remained stable across time (F_(3.66, 98.32)_ = 0.757, p > 0.05), **Fig. 3A**.

**Figure 3.**
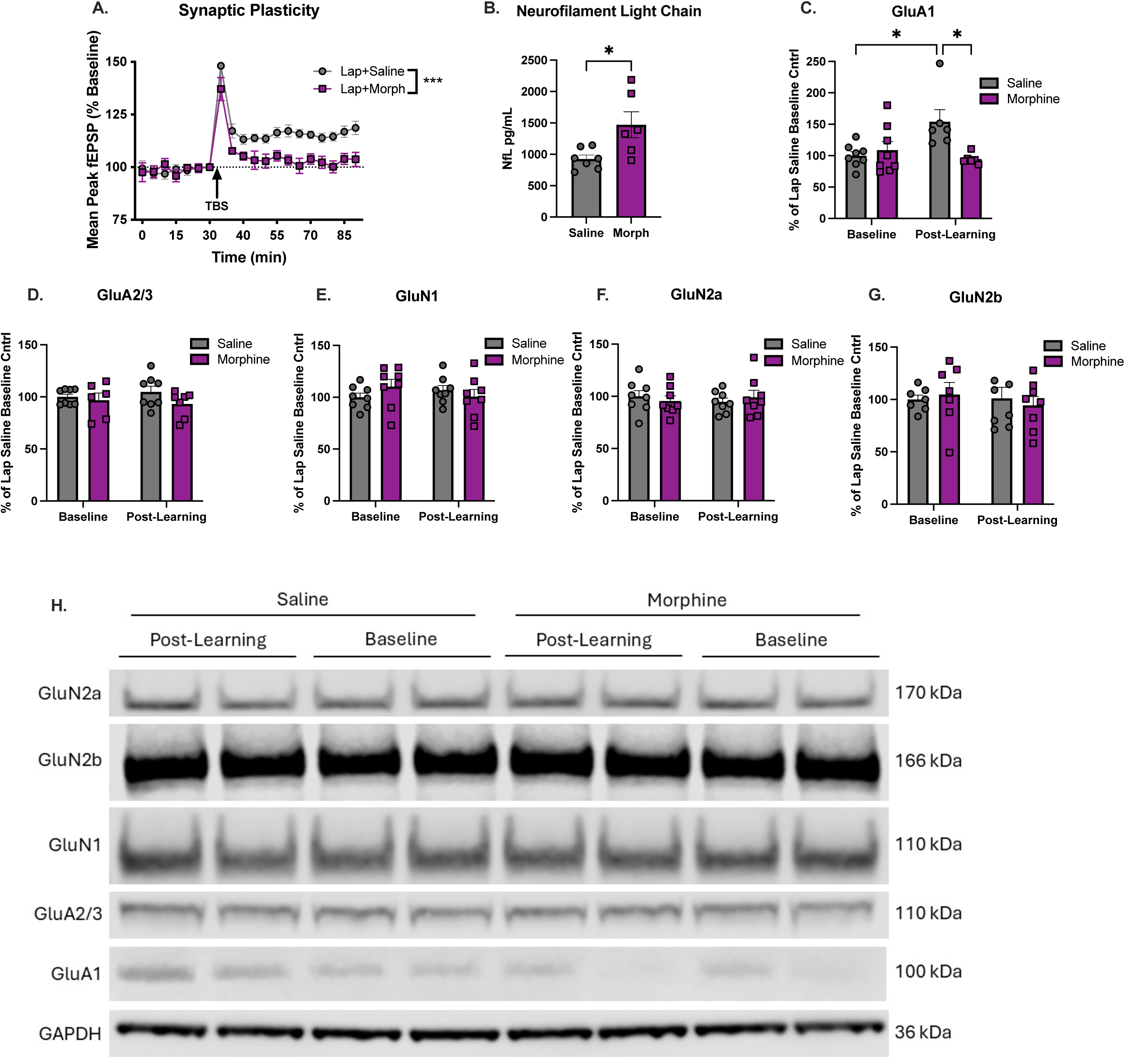
Synaptic plasticity, circulating neurofilament light chain, and glutamate receptor subunit expression. (A) Time course of field excitatory postsynaptic potential (fEPSP) response expressed as mean peak fEPSP (% baseline) following theta burst stimulation (TBS) in hippocampal slices from saline- and morphine-treated rats. ***p < 0.001 (B) Circulating neurofilament light chain (Nf-L) concentrations measured 30 days post-surgery in saline- and morphine-treated rats. Data are presented as mean ± SEM with individual data points shown. *p < 0.05. (C-G) Protein expression of glutamate receptor subunits GluA1 (C), GluA2/3 (D), GluN1 (E), GluN2a (F), and GluN2b (G) measured in hippocampal synaptoneurosomes under baseline conditions and following a learning experience, expressed as percent of laparotomy saline baseline control. (H) Representative immunoblots for glutamate receptor subunits and GAPDH corresponding to quantified data shown in panels C-G. Data are presented as mean ± SEM with individual data points shown. *p < 0.05.

#### Experiment 3

To determine whether long-lasting synaptic alterations were accompanied by broader neuroaxonal perturbation, we measured circulating Nf-L levels at day 30 after surgery. Circulating Nf-L levels at day 30 post-surgery were significantly greater in animals that received morphine than in those that received saline, indicating persistent neuroaxonal perturbation at this time point (t(11) = 2.72, p < 0.05; **Fig. 3B**).

Given that synaptic plasticity was clearly impaired four weeks after surgery in morphine-treated animals, and because glutamate receptors play a prominent role in synaptic plasticity due to their mediation of excitatory signals, we assessed protein expression of various glutamate receptor subunits. Expression was measured either at baseline or two hours following a learning experience, when memory consolidation is occurring. We found that GluA1 receptor expression was significantly altered by the combination of a learning experience and drug treatment (F_(1,24)_ = 7.95, p < 0.01), **Fig. 3C**. Specifically, having a learning experience significantly increased GluA1 expression in saline-treated rats compared to those that did not have a learning experience (p < 0.05), but this increase was significantly blunted in morphine-treated rats (p < 0.05). Expression levels of GluA2/3, GluN1, GluN2a, and GluN2b were not significantly altered with either learning experience or drug treatment (p > 0.05; **Fig. 3D-3G**).

#### Experiment 4

To begin to determine possible mechanistic targets, we measured 8-hydroxy-2’-deoxyguanosine (8-OHdG), a biomarker of DNA oxidation and readout of oxidative stress that is frequently elevated in chronic inflammatory conditions, especially with aging (Craft et al., 2012). We found that 8-OHdG was significantly elevated in the hippocampus of laparotomy and morphine-treated rats compared to saline-treated controls (t(12) = 3.40, p < 0.01), **Fig. 4A**. To assess whether this reflected a global alteration in cellular redox capacity, we measured NADPH, a ubiquitous cellular reducing equivalent, and observed no significant differences between the groups (t(12) = 0.56, p > 0.05), **Fig. 4B**. Together, these findings prompted further examination of mitochondrial function in the hippocampus, as mitochondrial oxidative phosphorylation is a major cellular source of reactive oxygen species and potential contributor to oxidative stress (Murphy, 2009).

**Figure 4.**
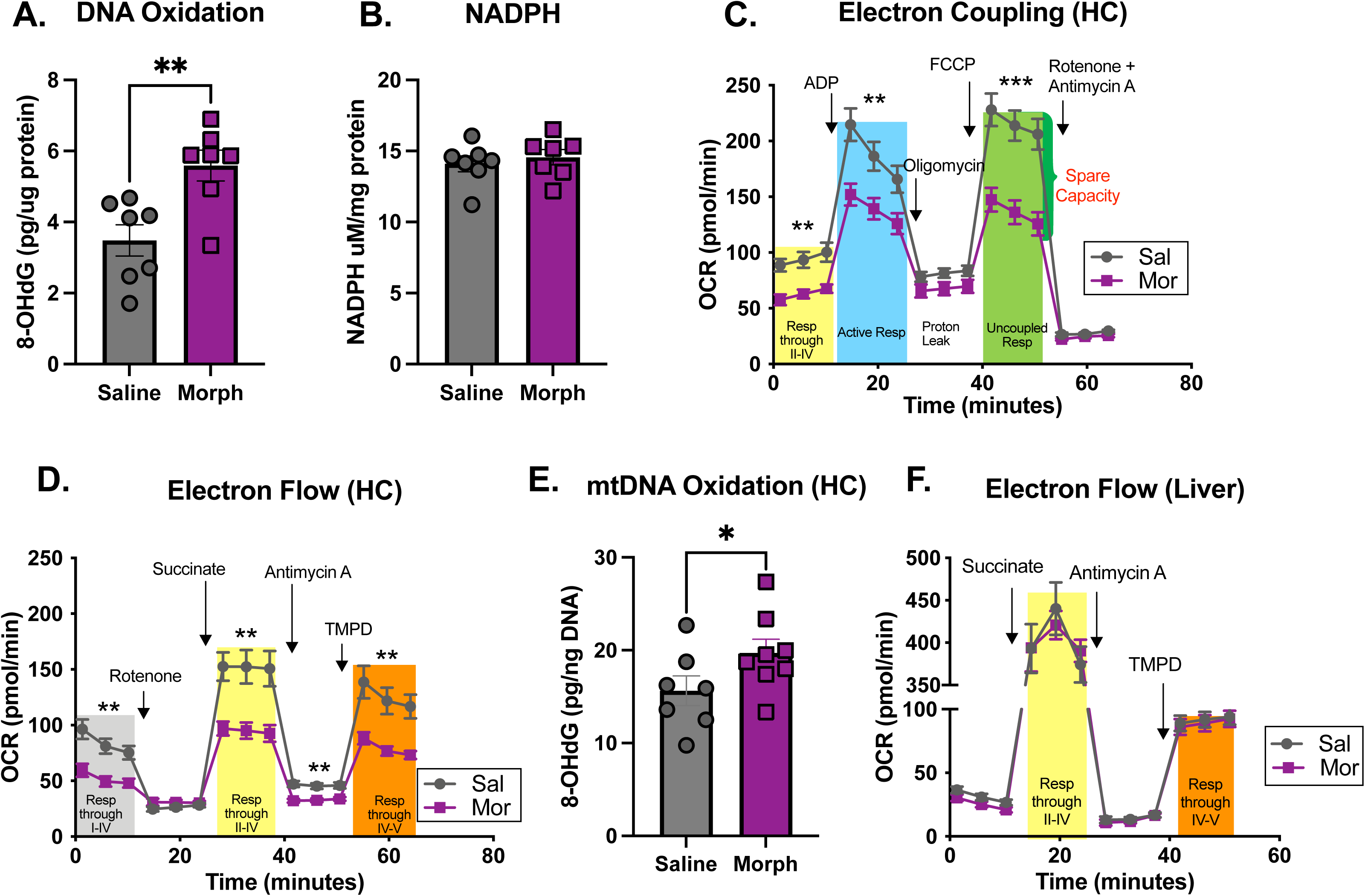
Oxidative stress markers and mitochondrial respiration in hippocampus and liver. (A) Levels of 8-hydroxy-2’deoxyguanosine (8-OHdG) measured in hippocampal tissue and expressed as pg/ug protein for saline- and morphine-treated rats. Data are presented as mean ± SEM with individual data points shown. **p < 0.01. (B) NADPH concentrations measured in hippocampal tissue and expressed as um/mg protein for each treatment group. Data are presented as mean ± SEM with individual data points shown. (C) Respiration of mitochondria isolated from fresh hippocampal tissue using the Seahorse electron coupling assay, showing oxygen consumption rate (OCR) over time following sequential injections of ADP, oligomycin, FCCP, and rotenone + antimycin A. Area under the curve was calculated for each phase. **p < 0.01, ***p < 0.001. (D) Respiration of mitochondria isolated from fresh hippocampal tissue using the Seahorse electron flow assay, showing OCR over time following sequential injections of rotenone, succinate, antimycin A and TMPD. Area under the curve was calculated for each phase. **p < 0.01. (E) Levels of 8-OHdG measured in mitochondrial DNA isolated from hippocampal tissue, expressed as pg/ng DNA. Data are presented as mean ± SEM with individual data points shown. *p < 0.05 (F) Respiration of mitochondria isolated from frozen liver tissue using the Seahorse electron flow assay, showing OCR over time following sequential injection of succinate, antimycin A, and TMPD. Area under the curve was calculated for each phase.

The electron coupling and electron flow Seahorse assays were run with fresh hippocampal tissue. Area under the curve (AUC) for each phase of both assays was calculated, followed by a Welch’s t-test to determine group differences. The electron coupling assay revealed significant impairments in mitochondrial respiration in morphine-treated rats compared to saline-treated controls. Specifically, morphine-treated rats exhibited reduced oxygen consumption rate (OCR) under basal conditions (t(8.34) = 3.83, p < 0.01), indicating diminished resting respiration. Following ADP injection, which stimulates active respiration, OCR remained significantly lower in the morphine-treated group (t(9.74) = 3.06, p < 0.01), indicative of impaired electron transport chain activity and the reduced ability of oxidative phosphorylation to meet high metabolic demands. Similarly, maximal respiratory capacity assessed after FCCP uncoupling was significantly reduced (t(9.98) = 4.58, p < 0.001), suggesting limited reserve respiratory capacity. In contrast, there were no differences following oligomycin treatment, which inhibits ATP synthase (t(10.62) = 1.89, p > 0.05), nor after combined rotenone & antimycin A treatment, which blocks electron transport-derived respiration (t(12.00) = 1.69, p > 0.05; **Fig. 4C**).

The electron flow assay also revealed significant impairments in mitochondrial respiration in morphine-treated rats compared to saline-treated controls. Specifically, morphine-treated rats exhibited reduced OCR under basal conditions (t(10.97) = 3.76, p < 0.01), indicating diminished uncoupled respiration. Following succinate injection, which stimulates Complex II-driven respiration, OCR remained significantly lower in the morphine-treated group (t(8.69) = 3.51, p < 0.01), suggesting impaired electron flow through electron chain complexes II through IV. OCR remained lower after antimycin A treatment, which inhibits Complex III and isolates upstream electron transfer (t(10.59) = 4.18, p < 0.001. Similarly, respiration following TMPD was significantly reduced (t(7.49) = 3.54, p < 0.01), indicating compromised Complex IV-dependent electron transfer. In contrast, there were no differences following rotenone treatment, which inhibits Complex I-mediated electron entry into the respiratory chain (t(12.81) = 1.36, p > 0.05; **Fig. 4D**).

Given the significant impairments observed in hippocampal mitochondrial respiration, we next assessed DNA oxidation specifically within hippocampal mitochondria. Mitochondrial DNA isolated from the hippocampus of morphine-treated rats exhibited elevated levels of 8-OHdG compared to saline-treated controls (t(12.66) = 1.87, p < 0.05; **Fig. 4E**), suggesting that mitochondrial DNA is particularly vulnerable to oxidative damage under these conditions.

To determine whether the mitochondrial deficits observed in the hippocampus were brain-specific or reflected more widespread peripheral dysfunction, we assessed OCR in mitochondria isolated from frozen liver tissue using the electron flow assay. No significant differences in uncoupled mitochondrial respiration were observed between morphine-treated rats and saline-treated controls at any phase of the assay including basal respiration (t(6.04) = 1.13, p > 0.05, succinate-stimulated respiration: (t(6.50) = 0.56, p > 0.05, antimycin A-inhibited respiration (t(5.16) = 0.48, p > 0.05, or TMPD-driven respiration (t(7.83) = 0.03, p > 0.05; **Fig. 4F**). These findings indicate that mitochondrial respiratory deficits are not evident in liver tissue and suggest mitochondrial dysfunction may be restricted to the hippocampus.

#### Experiment 5

To determine whether neuroinflammation directly contributes to the mitochondrial dysfunction observed at later stages of POCD, we pharmacologically inhibited TLR4 signaling in the brain at the time of surgery to suppress the initial inflammatory response and assessed oxidative stress and mitochondrial respiration four weeks later. Neuroinflammation is known to be elevated for at least two weeks in rats receiving both surgery and morphine (Muscat et al., 2021), raising the possibility that early inflammatory signaling contributes to persistent mitochondrial deficits.

Rats were assigned to one of three groups: 1) vehicle + surgery + saline, 2) vehicle + surgery + morphine, or 3) LPS-RS (TLR4 inhibitor) + surgery + morphine. Consistent with previous data (Fig. 1A), all rats exhibited a (∼2-5%) reduction in body weight during the morphine administration period compared to their pre-surgery weight, which they slowly regained over the next three weeks (F(1.51, 30.15) = 113.8, p < 0.0001), but there was no significant difference in weight between the rats that received saline, morphine, or combined morphine and LPS-RS treatment (F(2,20) = 0.045, p > 0.05; **Fig. 5A**).

**Figure 5.**
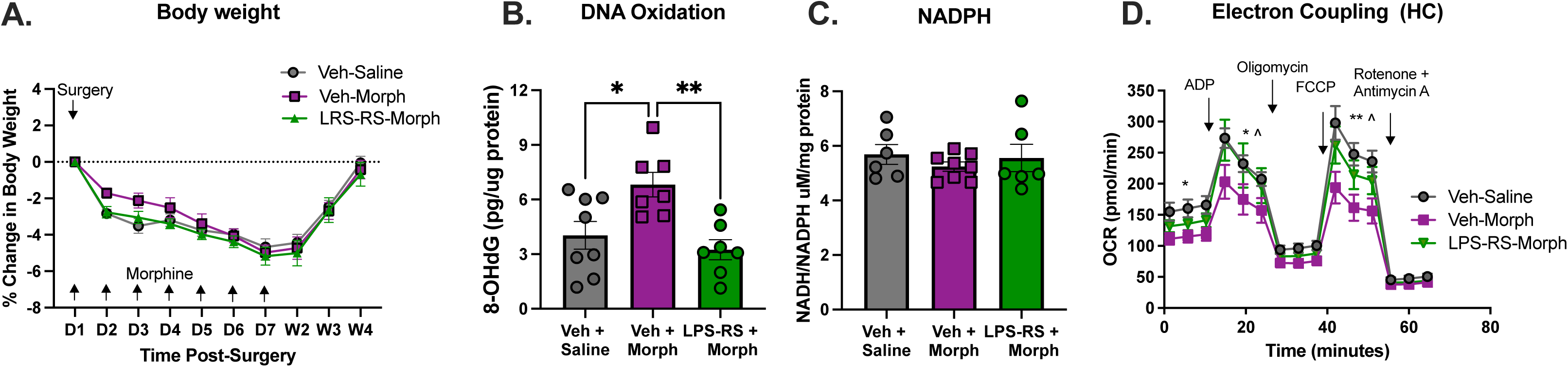
Effects of TLR4 inhibition on body weight, oxidative stress, and mitochondrial respiration following surgery and morphine. (A) Percent change in body weight relative to pre-surgery baseline across the post-surgical period in vehicle + saline- vehicle + morphine-, and LPS-RS + morphine-treated rats. Arrows indicate days of morphine administration. (B) Levels of 8-OHdG measured in hippocampal tissue and expressed as pg/ug protein for each treatment group. Data are presented as mean ± SEM with individual data points shown. *p < 0.05, **p < 0.01. (C) NADPH concentrations measured in hippocampal tissue and expressed as ug/mg protein for each treatment group. Data are presented as mean ± SEM with individual data points shown. (D) Respiration of mitochondria isolated from fresh hippocampal tissue using the Seahorse electron coupling assay, showing oxygen consumption rate (OCR) over time following sequential injections of ADP, oligomycin, FCCP, and rotenone + antimycin A for each treatment group. Area under the curve was calculated for each phase. *p < 0.05 between Veh-Saline and Veh-Morph, **p < 0.01 between Veh-Saline and Veh-Morph, ^p < 0.05 between LPS-RS-Morph and Veh-Morph.

One-way ANOVA revealed a significant group effect of DNA oxidation (F(2,19) = 7.45, p < 0.01; **Fig. 5B**). Consistent with earlier findings (Fig.4A), rats receiving vehicle + surgery + morphine exhibited significantly elevated levels of 8-OHdG compared to vehicle + surgery + saline controls (p < .05). In contrast, rats treated with the LPS-RS at the time of surgery and subsequently given morphine showed significantly reduced 8-OHdG levels compared to vehicle-treated morphine rats (p < 0.01) and were statistically indistinguishable from surgery + saline controls (p > 0.05). Similar to previous experiments (Fig. 4B), NADPH levels did not differ across any of the groups (F(2,17) = 0.48, p > 0.05; **Fig. 5C**).

A one-way ANOVA compared mitochondrial respiration of these three groups in the electron coupling Seahorse assay. There were significant group differences in mitochondrial respiration under basal conditions (F(2,19) = 3.97, p < 0.05), following ADP injection (F(2,18) = 3.63, p < 0.05), and after FCCP injection (F(2,19) = 5.57, p < 0.05; **Fig. 5D**). Post-hoc analyses revealed that vehicle + morphine-treated rats exhibited lower resting respiration compared to vehicle + saline-treated rats (p < 0.05), consistent with previous findings (Fig. 4C), but this was not attenuated with LPS-RS treatment (p > 0.05). However, oxidative phosphorylation and spare respiratory capacity in the vehicle + morphine group was significantly lower than both the vehicle + saline group (p < 0.05) and the LPS-RS + morphine group (p < 0.05) following ADP and FCCP injections. There were no differences between any of the groups following oligomycin treatment, which inhibits ATP synthase (F(2,19) = 2.20, p > 0.05), nor after combined rotenone & antimycin A treatment, which blocks electron transport-derived respiration (F(2,19) = 2.30, p > 0.05). The electron flow assay incurred some technical difficulties and therefore was unable to be analyzed. Collectively, these results suggest that inhibiting early central TLR4 inflammatory signaling can rescue morphine-induced mitochondrial dysfunction in the hippocampus.

#### Experiment 6

Given that mitochondrial dysfunction emerged as a prominent feature in later stages of POCD, we next set out to determine the extent to which rejuvenating mitochondria, with the novel drug SS-31, would ameliorate the memory deficit caused by surgery and morphine. First, we assessed body weight, to evaluate whether peripherally administered SS-31produced any systemic or adverse effects. Consistent with previous data, all rats exhibited a (∼2-5%) reduction in body weight during the morphine administration period compared to their pre-surgery weight, which they slowly regained over the next three weeks (F(3.672, 47.74) = 105.7, p < 0.0001), but there was no significant difference in weight between the rats that received SS-31 and saline treatment at week 4 (F(1,13) = 0.077, p > 0.05), **Fig. 6A**.

**Figure 6.**
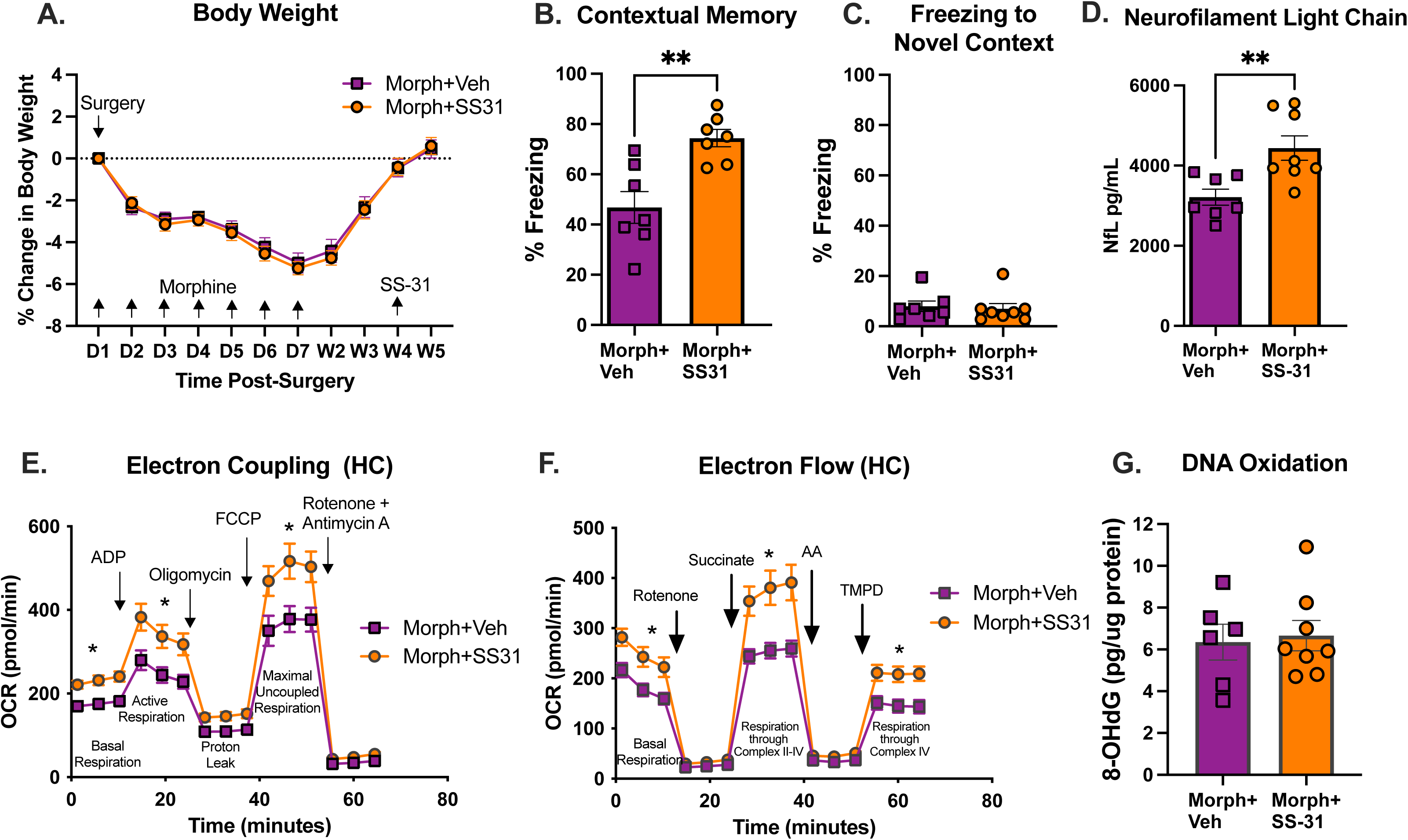
Effects of SS-31 on behavior, neuroaxonal biomarkers, mitochondrial respiration, and oxidative stress following surgery and morphine. (A) Percent change in body weight relative to pre-surgery baseline across the post-surgical period in morphine + vehicle and morphine + SS031-treated rats. Arrows indicated days of morphine administration and SS-31 treatment. (B) Freezing behavior expressed as percent freezing during the contextual fear memory tests in morphine + vehicle- and morphine + SS-31-treated rats. Data are presented as mean ± SEM with individual data points shown. **p < 0.01. (C) Freezing behavior expressed as percent freezing in a novel context for each treatment group. Data are presented as mean ± SEM with individual data points shown. (D) Circulating neurofilament light chain (Nf-L) concentrations measured 30 days post-surgery in morphine + vehicle and morphine + SS031-treated rats. Data are presented as mean ± SEM with individual data points shown. **p < 0.01. (E) Respiration of mitochondria isolated from fresh hippocampal tissue using the Seahorse electron coupling assay, showing OCR over time following sequential injections of ADP, oligomycin, FCCP, and rotenone + antimycin A for each treatment group. Area under the curve was calculated for each phase. *p < 0.05. (F) Respiration of mitochondria isolated from fresh hippocampal tissue using the Seahorse electron flow assay, showing OCR over time following sequential injection of rotenone, succinate, antimycin A and TMPD for each treatment group. Area under the curve was calculated for each phase. *p < 0.05. (G) Levels of 8-OHdG measured in hippocampal tissue and expressed as pg/ug protein for each treatment group. Data are presented as mean ± SEM with individual data points shown.

In prior reports from our laboratory, contextual fear memory in this paradigm typically yields freezing levels of approximately 65-70% in unimpaired animals, whereas morphine-treatment produces a robust reduction in freezing behavior (∼40-45%) (Muscat et al., 2023; Muscat et al., 2021). Consistent with our previous studies, morphine-treated rats exhibited impaired contextual freezing (46%), while combined morphine + SS-31-treatment significantly ameliorated this deficit, increasing freezing to 74% (t(12) = 3.823, p < 0.01; **Fig. 6B**). In contrast, freezing behavior in a novel context remained low (∼5%) and did not differ significantly between groups (t(13) = 0.339, p > 0.05; **Fig. 6C**). These data indicate that SS-31 treatment administered during advanced stages of POCD is sufficient to rescue the ability to form new memories.

To assess whether SS-31 treatment altered neuroaxonal biomarkers at this time point, circulating Nf-L levels were measured 30 days post-surgery. Circulating Nf-L levels following surgery + morphine were significantly greater in animals that received SS-31 than in those that received saline at day 30 (t(11.78) = 3.36, p < 0.01; **Fig. 6D**). When considered alongside the behavioral findings, this dissociation indicates that elevated Nf-L occurred despite functional recovery.

Given the robust rescue of hippocampal-dependent memory, we next examined whether SS-31 ameliorated the mitochondrial deficits observed in surgery + morphine-treated animals.

The electron coupling assay showed significant amelioration of mitochondrial dysfunction in animals treated with SS-31 at the 30-day timepoint. Specifically, compared to animals treated with saline, SS-31-treated rats exhibited greater resting mitochondrial respiration (t(13) = 2.57, p < 0.05), greater ATP-linked respiration (t(12.03) = 2.08, p < 0.05), and greater maximal respiration (t(12.46) = 1.99, p < 0.05). As expected, there were no differences following oligomycin treatment, which inhibits ATP synthase (t(12.00) = 1.75, p > 0.05), nor after combined rotenone & antimycin A treatment, which blocks electron transport-derived respiration (t(12.91) = 1.53, p > 0.05; **Fig. 6E**).

The electron flow assay similarly revealed a significant amelioration of mitochondrial respiration in SS-31-treated rats compared to those treated with saline. Specifically, SS-31-treated rats exhibited increased uncoupled basal respiration (t(10.92) = 2.10, p < 0.05), increased Complex II-driven respiration (t(9.08) = 2.43, p < 0.05), and increased Complex IV-dependent respiration (t(11.85) = 2.33, p < 0.05). As expected, there were no differences following rotenone treatment, (t(10.61) = 1.71, p > 0.05) or antimycin A (t(10.68) = 1.86, p > 0.05; **Fig. 6F**).

Finally, we assessed DNA oxidation in the hippocampus and found that SS-31 did not reduce surgery + morphine-induced levels of 8-OHdG (t(10.84) = 0.27, p > 0.05; **Fig. 6G)**. Collectively, the results suggest that SS-31 holds promise as a therapeutic intervention to minimize the negative consequences of established POCD.

## Discussion

The present study demonstrates that perioperative morphine administration produces long-lasting impairments in hippocampal-dependent memory that persist weeks after surgery and are accompanied by selective synaptic, mitochondrial, and oxidative abnormalities in the hippocampus. Across a series of complementary experiments, we show that these cognitive deficits are not attributable to gross changes in dendritic architecture or systemic illness, but instead coincide with reduced dendritic spine maturity, impaired long-term synaptic plasticity, elevated circulating Nf-L, altered experience-dependent AMPA receptor expression, elevated oxidative stress, and sustained mitochondrial respiratory dysfunction. Importantly, pharmacological interventions targeting either early neuroinflammatory signaling or later mitochondrial function partially normalized these abnormalities, revealing a dissociation between persistent oxidative damage and recoverable mitochondrial bioenergetics.

Despite transient postoperative weight loss in all animals, body weight recovered fully by three weeks post-surgery and did not differ consistently across treatment groups, indicating that long-term cognitive and cellular alterations are unlikely to reflect nonspecific sickness behavior. Likewise, morphine treatment did not alter overall dendritic arborization of CA1 pyramidal neurons, suggesting that persistent cognitive deficits are not driven by large-scale structural degeneration. Instead, morphine selectively reduced dendritic spine populations associated with synaptic stability and plasticity, consistent with what others have reported (Robinson & Kolb, 1999, 2004). We found mushroom spines in both apical and basal compartments and thin spines in basal dendrites were particularly diminished. Given the established role of these spine subtypes in maintaining synaptic strength and supporting long-term potentiation (Borczyk, Sliwinska, Caly, Bernas, & Radwanska, 2019; Bourne & Harris, 2007; Runge, Cardoso, & de Chevigny, 2020), these changes are consistent with a functional weakening of synaptic connectivity rather than synapse loss.

Consistent with this interpretation, morphine-treated animals exhibited a marked impairment in late-phase LTP several weeks after surgery, despite intact baseline synaptic transmission. This pattern mirrors prior behavioral findings demonstrating preserved short-term memory alongside disrupted long-term memory (Barrientos et al., 2006; Muscat et al., 2021), and points to a failure of activity-dependent synaptic consolidation mechanisms (Chapman et al., 2010). Supporting this conclusion, learning-induced upregulation of the AMPA receptor subunit GluA1 was blunted in morphine-treated rats, whereas other glutamatergic receptor subunits were not detectably altered at the sampled time point. This pattern may reflect a selective vulnerability of GluA1-dependent plasticity mechanisms. However, it is also possible that regulation of other subunits was missed due to differences in post-activity kinetics.

Glutamate receptor subunits can exhibit distinct temporal dynamics in trafficking, local translation, and surface redistribution following synaptic activity and LTP induction, with some changes occurring rapidly (minutes) and others emerging later (tens of minutes to hours) (Cercato et al., 2016; Dupuis et al., 2014; Newpher & Ehlers, 2008). Future work sampling multiple post-learning time points and separating surface versus total receptor pools will help clarify whether morphine-induced POCD broadly perturbs glutamate receptor regulation or preferentially impairs GluA1-dependent consolidation.

In parallel with synaptic dysfunction, animals receiving surgery and morphine exhibited elevated circulating neurofilament light chain (Nf-L) at 30 days post-surgery, consistent with persistent neuroaxonal stress. Nf-L is a major axonal cytoskeletal protein whose release into circulation is widely used as a sensitive marker of neuroaxonal injury and cytoskeletal turnover, and may also reflect remodeling processes rather than irreversible neurodegeneration per se (Gafson et al., 2020; Pekny et al., 2021). Thus, elevated Nf-L in this context likely reflects sustained axonal strain or increased cytoskeletal turnover associated with prolonged synaptic dysfunction and mitochondrial bioenergetic impairment.

A central finding of this study is that these synaptic and cognitive impairments are accompanied by sustained mitochondrial dysfunction and oxidative stress localized to the hippocampus. Morphine-treated animals exhibited elevated levels of 8-OHdG, indicating increased DNA oxidation (Cooke, Evans, Dizdaroglu, & Lunec, 2003; Valavanidis, Vlachogianni, & Fiotakis, 2009), without alterations in global cellular reducing capacity as indexed by NADPH (Pollak, Dolle, & Ziegler, 2007). Seahorse analyses revealed pronounced impairments in mitochondrial respiration, including reduced basal respiration, diminished ATP-linked oxidative phosphorylation, and limited spare respiratory capacity. These deficits spanned multiple components of the electron transport chain, including Complex II- and Complex IV-dependent respiration, and were accompanied by elevated mitochondrial DNA oxidation. Importantly, comparable mitochondrial impairments were not observed in liver tissue, indicating that these effects are regionally specific rather than reflective of systemic metabolic dysfunction.

Mechanistically, our findings further support a critical role for early neuroinflammatory signaling in initiating long-lasting mitochondrial dysfunction. Pharmacological inhibition of TLR4 signaling at the time of surgery attenuated hippocampal DNA oxidation and partially restored mitochondrial capacity weeks later, indicating that early inflammatory processes contribute to the persistence of mitochondrial deficits. Importantly, in a prior POCD study using the same dose and timing of the TLR4 inhibitor LPS-RS, we demonstrated that blocking TLR4 signaling at the time of surgery completely ameliorated the long-term cognitive deficit induced by surgery and morphine (Muscat et al., 2023). Together with the present data, these findings suggest that early TLR4-dependent neuroinflammatory signaling is sufficient to drive both cognitive impairment and downstream mitochondrial dysfunction. The incomplete normalization of basal respiration observed here further suggests that while early neuroinflammation signaling is a key initiating factor, additional mechanisms, possibly including sustained metabolic or opioid-mediated effects, may contribute to the maintenance of mitochondrial impairment at later time points as others have suggested using various models (Feng et al., 2008; Reymond, Vujic, Schvartz, & Sanchez, 2022; Sun, Torices, Osborne, & Toborek, 2025). Future studies should examine the contribution of these additional factors in our POCD model.

Crucially, targeted mitochondrial rejuvenation with SS-31 significantly improved mitochondrial respiratory function and rescued hippocampal-dependent memory despite persistently elevated DNA oxidation. SS-31-treated animals exhibited enhanced basal respiration, increased ATP-linked oxidative phosphorylation, and greater maximal respiratory capacity in the electron coupling assay, as well as improved Complex II- and Complex IV-dependent respiration in the electron flow assay. These improvements were observed at a time point when hippocampal 8-OHdG levels remained elevated and were indistinguishable from those in surgery + morphine-treated animals receiving saline. This dissociation indicates that restoration of mitochondrial bioenergetic efficiency can occur independently of detectable reductions in accumulated DNA oxidation, at least in the short term. Whether sustained oxidative DNA damage ultimately compromises longer-term synaptic or cognitive outcomes remains an important question for future investigation.

These findings are consistent with the proposed mechanism of SS-31, which stabilizes cardiolipin-cytochrome c interactions and improves electron transport efficiency rather than acting as a direct antioxidant (Birk et al., 2013; Chavez et al., 2020; Szeto, 2014). Thus, SS-31 may rescue cognitive function through optimization of mitochondrial performance even in the presence of persistent oxidative stress markers.

At the behavioral level, SS-31 robustly ameliorated contextual fear memory deficits while leaving freezing in a novel context unchanged, indicating a selective rescue of hippocampal-dependent memory rather than generalized behavioral activation. Together with the mitochondrial findings, these results support a model in which mitochondrial dysfunction is a proximal driver of long-term cognitive impairment following surgery and morphine exposure, and that improving mitochondrial efficiency is sufficient to restore memory performance even when oxidative damage remains unresolved.

Interestingly, SS-31 treatment further increased circulating Nf-L levels despite robust rescue of hippocampal-dependent memory and normalization of mitochondrial respiratory function. While elevated Nf-L is often interpreted as a marker of axonal injury, increasing evidence indicates that Nf-L can also reflect heightened neuroaxonal turnover and cytoskeletal remodeling during periods of recovery or increased plasticity (Gafson et al., 2020; Pekny et al., 2021). In this context, elevated Nf-L following SS-31 treatment may reflect increased neuroaxonal turnover or remodeling accompanying restored synaptic function, rather than exacerbated neurodegeneration. This interpretation is supported by the dissociation observed here between Nf-L levels and functional outcome, as SS-31-treated animals exhibited marked improvements in memory performance and mitochondrial bioenergetics despite elevated Nf-L.

Several limitations warrant consideration. Normal freezing levels were inferred from prior studies using the same behavioral paradigm (Muscat et al., 2024; Muscat et al., 2023; Muscat et al., 2021). However, the consistency of these benchmarks across experiments strengthens their interpretive value. Additionally, while SS-31 restored mitochondrial function at the measured time point, it remains unknown whether longer treatment durations would ultimately reduce oxidative damage, or if a longer observation window would reveal greater oxidative damage and worsening cognitive function. Future studies should examine the temporal relationship between oxidative stress, mitochondrial dysfunction, and synaptic recovery, as well as determine whether combined antioxidant and mitochondrial therapies provide additive benefit. Future studies incorporating longitudinal Nf-L measurements and direct assessment of axonal structure will be necessary to distinguish injury-related release from plasticity-associated neurofilament turnover.

In summary, this work identifies sustained hippocampal mitochondrial dysfunction as a key mechanism underlying long-term cognitive deficits following surgery and morphine exposure. By demonstrating that mitochondrial bioenergetic function, and cognitive performance, can be restored independently of persistent DNA oxidation, these findings refine current models of POCD and highlight mitochondria as a viable therapeutic target even after oxidative damage has occurred.

## Acknowledgements

This work was supported in part by RF1-AG028271 from the National Institute on Aging (to RMB); R01HL166520 (to KKB) and T32HL134616 (to HFS and DWK) from the National Heart, Lung, and Blood Institute.

## Notes

### Competing Interest Statement

The authors have declared no competing interest.

